# Structured interactions explain the absence of keystone species in synthetic microcosms

**DOI:** 10.1101/2025.03.31.646297

**Authors:** Sivan Pearl Mizrahi, Hyunseok Lee, Akshit Goyal, Erik Owen, Jeff Gore

## Abstract

In complex ecosystems, the loss of certain species can trigger a cascade of secondary extinctions and invasions. However, our understanding of the prevalence of these critical “keystone” species and the factors influencing their emergence remains limited. To address these questions, we experimentally assembled microcosms from 16 marine bacterial species and found that multiple extinctions and invasions were exceedingly rare upon removal of a species from the initial inoculation. This was true across eight different environments with either simple carbon sources (e.g., glucose) and more complex ones (e.g. glycogen). By employing a generalized Lotka-Volterra model, we could reproduce these results when interspecies interactions followed a hierarchical pattern, wherein species impacted strongly by one species were also more likely to experience strong impacts from others. Such a pattern naturally emerges due to observed variation in carrying capacities and growth rates. Furthermore, using both statistical inference and spent media experiments, we inferred interspecies interaction strengths and found them consistent with structured interactions. Our results suggest that the natural emergence of structured interactions may provide community resilience to extinctions.

## Introduction

A long-standing question in ecology is how an ecosystem reacts to the removal of a resident species. While the extinction of one species may leave a minimal impact on the community structure, the extinction of another species may lead to drastic changes, including cascades of extinction and the entry of invasive species. In his pioneering work Robert Paine coined the term “keystone” species^1^, describing the starfish predator *Pisaster ochraceus* whose removal from the ecosystem led to a dramatic decrease in biodiversity. Other early identified keystone species included sea otters, whose role as an apex predator regulated the density of sea urchins and maintained kelp forests^2^. However, use of the term has expanded to denote a species that exerts a substantial impact on a community or ecosystem, disproportionate to its abundance^3^. A plethora of theoretical studies suggest that keystone species should be prevalent in natural ecosystems^4–8^. These studies focus on different properties of the species-interaction network to predict “keystoneness”. For example, highly connected species are believed to be more likely to serve as keystone species. Understanding the prevalence of keystone species within a community is essential both for conservation efforts and for community engineering.

Environmental factors such as resource availability are thought to be important in determining the frequency of keystone species. The interactions between different species can change in different environments, such that changing the resources can cause biotic-interactions to shift^3,9,10^. Thus, certain species can function as a keystone in one environment but not in another^10,11^. In microbial communities, specifically, the metabolic abilities of certain species could dictate their role as keystone species, for instance, when they are performing a “key function” such as being the only efficient degraders of a recalcitrant substrate^12^. Changing the resources available within an environment could therefore alter the identity and even the presence of keystone species.

A growing body of work suggests microbial communities are also likely to include keystone species^13–16^. Microbial communities offer practical tools to study the resilience of a system to species removal. Current microbial studies are fueled by high-throughput sequencing studies of different microbiomes and their sampling through time and space. Such serial data were used to infer the interactions among species and predict keystone species from network properties^15^. Utilizing synthetic microbial communities to perform community “drop-out” experiments, several studies have explored the impact that species removal has on a community^17–20^. Carlström, *et al*,

Venturelli *et al* and Weiss *et al* have identified certain species whose ‘dropping-out’ significantly impact community structure (though not focusing specifically on secondary extinctions or invasions)^10,17,20^. Niu *et al*. studied a synthetic seven-species maize-root community and identified *Enterobacter cloacae* as a keystone species whose removal from the initial inoculation of the community results in five secondary extinctions and the dominance of the remaining species^19^. These studies exemplify the use of synthetic microbial communities as a practical tool to identify species with a strong impact on community structure. Despite the identification of particular communities with a keystone species, the prevalence of keystone species across different environments remains unclear.

Here, we examined how frequently species removal from the community leads to multiple secondary invasions and extinctions, and how the frequency of such cascades changes under varying growth conditions. We comprehensively assessed the prevalence of keystone species across a range of resources in bacterial microcosms, utilizing a synthetic marine microbial species pool to systematically explore the impact of each species’ removal. We assembled communities from the same pool of 16 species in eight distinct conditions, resulting in communities with varying species richness. Despite theoretical predictions suggesting that keystone species should be common, we find a complete absence of keystone species across all eight environmental conditions. Modifying the generalized Lotka-Volterra (gLV) model to include a hierarchical structure of interspecies interactions, expected to naturally arise from carrying capacity variation, recapitulates the lack of keystone species. We provide experimental evidence that species interactions in our communities indeed have such a structure. Together, our findings indicate that keystone species may be less common than previously assumed, partly owing to the natural emergence of structured interspecies interactions.

## Materials and Methods

### Species, media, and carbon sources

We have used in this work 16 isolates from the PRiME project of the Simons Foundation^24,25^, detailed in Table S1.

Marine broth (MB) (Difco 2261) supports the growth of all 16 species used in this work. MB was used for streaking, plating, and growth of starter cultures.

All experiments were carried in minimal media based on a recipe from Amaranth *et al*^26^ (detailed in supplementary methods), supplemented with different carbon sources listed in Table S2.

### EKO experiments

Colonies from all 16 isolates grown on MB-agar plates were picked and placed into culture tubes with 3ml MB, grown at room temperature with shaking at 300rpm for two days. ODs were measured and volumes equal to 0.006 OD_600_ were taken from each isolate. 17 assemblies were prepared: one inoculation with all 16 isolates and 16 with each isolates removed. These were diluted 1:80 into the respective minimal media with the relevant carbon source. For each community and media combinations there were either triplicate or duplicate samples. The growth dilution cycles were carried in 1ml deep 96-well plates (Eppendorf EP951032701) filled with 400ul, sealed with membrane (Excel Scientific AeraSeal sterile). For each cycle the plates were grown at 25°C, shaking at 300rpm for 48hr. Using a multichannel pipettor (Viaflo 96, Integra Biosciences, Hudson, USA) plates were mixed and 10uL were transferred to 390uL of fresh matched media, mixed, sealed, and incubated again. Following transfer, 100ul remaining from the previous cycle were taken to a 96 well cell culture plate (VWR 62406-081) and OD_600_ was measured using a Tecan infinite 200 pro multiplate reader. The 7^th^ cycle was frozen at -80°C and later taken for DNA extraction and 16S-amplicon sequencing (Supplementary methods).

### Inferring EKO impacts

We developed a threshold-based method to assess the influence of species EKO on community structure, accounting for variability across replicates and species whose relative abundances fluctuated independently of the specific EKO.

Two approaches were used to infer whether the knockout of species *i* impacts species *j*:

#### Threshold based

For a given threshold *t*, species *i* is considered occurring in a sample if its frequency exceeds *t*. Each community consists of replicates grown in the same media. We denote *O*^*i,m*^ to be the mean of occurrences of species *i* in condition *m* over all replicates. Species *i* is considered present in condition *m* if it occurs in the majority of replicates for that condition (*O*^*i,m*^*>0*.*5*^)^.

Species *i* is considered invading in condition *m* if the difference in species *i*’s occurrences between the EKO and the full community exceeds 0.5: *O*^*i,m*^*-O*^*i,full*^ >0.5 and extinct if its<0.5.

We have found a subset of species display high variability in their occurrences across all samples of a certain media. Species for which the mean occurrences between all samples in the given media (all EKOs and the full community) ranged between 0.2-0.8 considered inconsistent. We are assuming that in most EKO each species *i* is likely to have similar frequency as in the full community, while only in certain EKOs of impactful species it would differ. Large variability is unexpected and indeed was found to be unrelated to the background community in most cases. as their occurrences general seemed to be independent of the background community. For such species, we refined our analysis by comparing their mean frequencies in EKO communities to their overall mean frequency across all samples, considering only species whose mean frequency +/-std in the EKO are lower than the mean frequency -\+std over all samples as extinct\invading, respectively.

We have run this analysis on 13 thresholds: [0.001, 0.0015,0.002…0.007] and species were considered to experience secondary impacts if they were impacted in more than 6 of these thresholds.

This method tries to identify invasions and extinctions and thus ignores many differential abundances scenarios. Due to our relatively small synthetic communities’ setup and small number of samples for each condition (<=3), many of the assumptions made in microbiome analysis frameworks that are used for differential abundance analysis are not suitable.

#### ANOVA based

We also conducted ANOVA on centered log-ratio transformed frequencies to detect species that exhibited significantly different means in specific EKO conditions compared to others. While we acknowledge that the normality assumption of ANOVA may not be met, non-parametric Kruskal-Wallis tests did not detect significant impacts, so we present the ANOVA results instead.

### Model

We use the following equation to study ecological KO in the presence of carrying capacity:

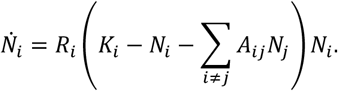

Here *N*_*i*_ is the population size of *i*-th species, *K*_*i*_ is the carrying capacity of *i*-th species, and *A*_*ij*_ is the interaction between species *i* (affected) and *j* (affecter). More details on parameters and simulations are in the Supplementary data.

## Results

To study the sensitivity of a community to single species removal we need to consider both the community members and the environment, which plays a significant part in shaping the community structure. Species have different properties in different environments (e.g., different growth rates, carrying capacities, etc.), and interspecies interactions between species can also change between different environments. We chose to study a pool consisting of 16 bacterial species isolated from marine coastal waters that are representative of 5 major families (*Flavobacteriaceae, Rhodobacteraceae, Oceanospirillaceae, Alteromonadaceae* and *Vibrionaceae)* that could be differentiated by 16S amplicon-sequencing (Table S1). In addition, we studied eight different environments that differ from one another by their carbon source (Table S2). The carbon sources we used range in complexity from being very simple molecules (such as acetate and glucose) that are accessible to most species to long polymers (such as alginate and glycogen) that require specialized enzymes to break down. We also included an environment composed of many different carbon sources e.g., amino acids and sugar sources (denoted MCS, for multiple carbon sources). When grown by themselves in monoculture the species display large variability in growth capabilities, with less than half of the species displaying significant growth on sucrose but with all species growing well in the MCS media (Fig. 1A). The wide range of environments, together with the associated variation in species’ ability to grow on the provided carbon source, ensure that our assembled communities have a variety of different interactions and dependencies.

**Figure 1.**
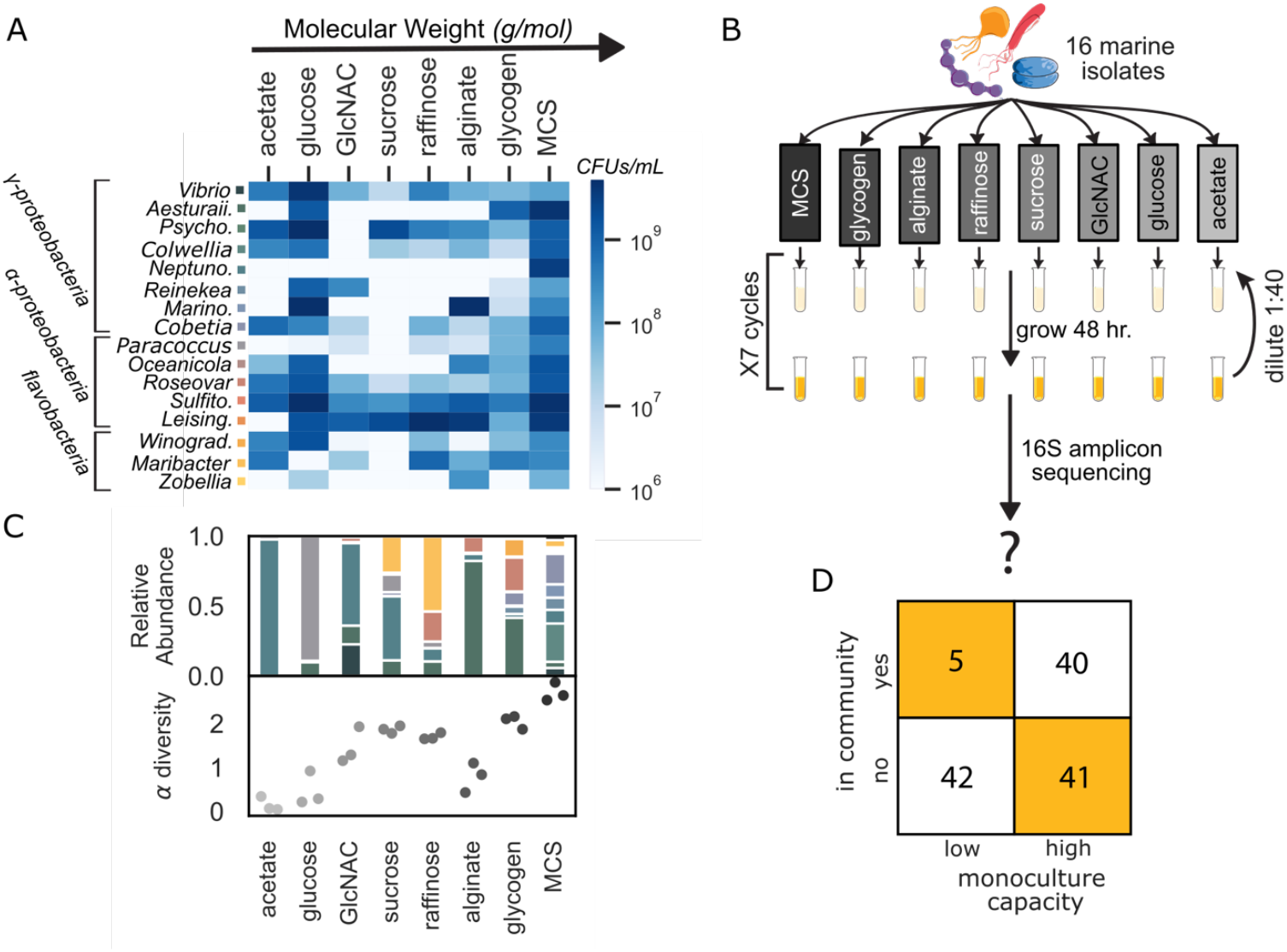
Synthetic microcosmos of 16 marine bacteria assembled on 8 different carbon sources differ in biodiversity and harbor multiple interactions. **A**. Heatmap of the capacities (CFUs/mL) of each species’ monoculture at the end of four growth-dilution cycles. Each row designates a different species, ordered according to their family phylogenetic classification **B**. Experiment layout: 16 marine bacterial species are inoculated in samples with minimal media with different carbon sources and are grown in 7 growth-dilution cycles of 1:40 every 2 days. At the end of the 7^th^ cycle the samples are processed and sent to 16S-amplicon sequencing to probe the community structure. **C**. Biodiversity of the different communities. Top panel: the mean relative abundance of each species in each community over triplicates. Relative abundance is calculated for each sample: 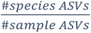Colors of species are same as in A. Bottom panel: the alpha diversity of each sample, using the Shannon index. **D**. Growth capability as monoculture is not necessarily indicative of survival in the community. Count table of all species capacities in monoculture (below or above 10^7^ CFUs/mL) and their appearance in the community (>0.004% relative abundance).

### Communities assembled on different carbon sources harbor multiple interactions

We first assembled communities by inoculating media of the different environments with all 16 species, which we refer to as “full” communities. The full communities were grown in triplicate and diluted 1:40 every 2 days for 7 cycles (Fig. 1B). 16S-amplicon sequencing revealed that the community composition of the full communities varied with environment, with greater biodiversity in the more complex carbon sources (either with higher molecular weight or being MCS with many different carbon sources). For example, while the acetate environment was inhabited by one dominant species (and one other species with very low frequency that crosses our threshold of 0.4% in only one of the triplicates), the MCS environment harbored 10 abundant species without distinct dominance (Fig. 1C). Many of the species that were unable to grow well in monoculture were nonetheless able to coexist in the community (Fig 1D). At the same time, many of the species that grew well in monocultures were unable to survive in the community (Fig 1D). Both observations suggest that both beneficial and competitive interactions are widespread within our communities.

### Lack of keystone species in experimental marine communities

We next developed an experimental framework to assess the likelihood of secondary extinctions and invasions of community members resulting from the removal of a particular species from the community. Since removing a specific species from the assembled community is technically challenging, we used an alternative approach we termed “Ecological Knock-Outs” (EKOs) in which we assembled distinct communities by systematically removing a specific species from the initial inoculum used during community assembly (sometimes called drop-out experiments). Species that were present in the full community but were not present in the assembled EKO community constituted secondary extinctions, while species that were not in the full community yet were in the EKO constituted secondary invasions. To determine which species are secondarily impacted and address noise in our dataset, we used thresholds and a majority-decision rule based on replicates. A species is considered impacted if the majority of replicates fall above or below multiple thresholds compared to the majority of replicates in the full community (see Methods). A keystone species would be a species whose EKO leads to multiple secondary impacts (both invasions and extinctions).

To characterize the frequency of keystone species we experimentally performed knockouts of each of the sixteen species in each of the eight environments. For example, in glycogen the full community contained a moderately diverse set of seven species that were able to coexist together (Fig 1C and Fig 2A). As expected, experimentally removing each of the nine species that did not survive in the full community led to no change in their respective EKO communities. However, among the seven species that were present within the full community, we found only two examples in which species removal led to secondary impacts on the remaining community. In particular, removal of the *Neptuno*. species led to a secondary invasion of the *Marino*. species, in that the

**Figure 2.**
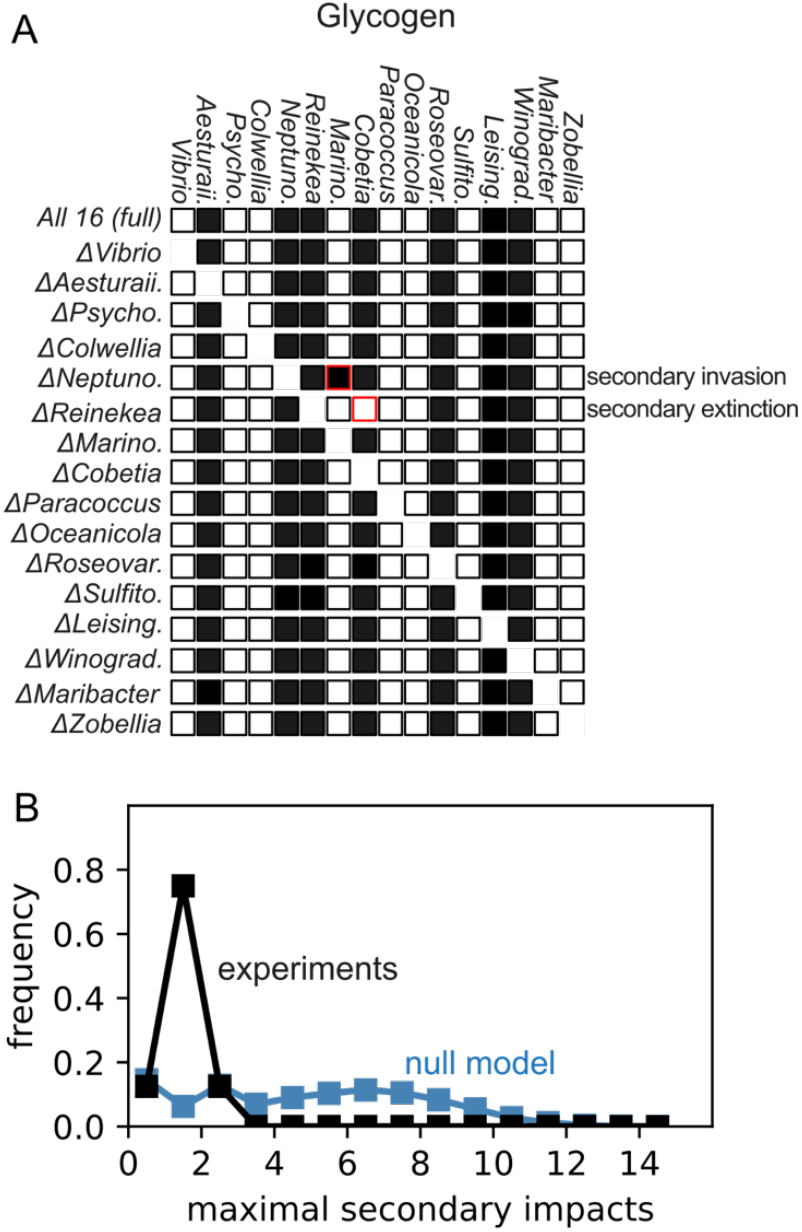
Lack of keystone species in experimental marine microcosms assembled in eight different carbon sources. **A**. Schematic illustration of species survival of the full and all EKO communities, grown on glycogen as the carbon source. Each row represents a community, and each column represents a species. Filled and empty squares indicate survival and extinction, respectively. A red frame indicates a secondary impact where this species’ presence differ from that in the full community. Here, the EKO of Neptuno. led to the secondary invasion of Marino and the EKO of Reinekea led to the secondary extinction of Cobetia. **B**. Histogram of the maximum secondary impacts per EKO of all EKOs in all eight environments. This is a measure that reflects the EKO with most secondary impacts for a given ecosystem (carbon source/simulation), which describe the “keystone-ness”. The null model simulation is in blue, and the experimental data is in black.

*Marino*. species was not present in the full community but was able to survive in the community when the *Neptuno*. was excluded. In addition, removal of the *Reinekea* species led to secondary extinction of the *Cobetia* species (Fig 2A). Despite the presence of secondary impacts upon removal of two of the species, neither qualifies as a keystone species as the secondary impacts were modest and did not entail a cascade of secondary impacts and associated wholesale changes to the community. Our ecological knockout experiments therefore indicate that there was no keystone species in the glycogen environment.

Across all eight environments, the overwhelming majority of ecological knockouts (115 out of 128) resulted in no secondary impacts, with species composition remaining unchanged before and after species removal. Among the remaining 13 EKOs, 12 led to a single secondary impact, 11 of which were invasions, corresponding to the replacement of the removed species by another. The only secondary extinction observed was the one noted previously in glycogen. The most impactful EKO belonged to *Neptuno*., the dominant species on acetate, whose removal led to the invasion of two species (Supplementary Figure 1). Given the potential for up to 15 secondary impacts, even *Neptuno* likely does not qualify as a keystone species in acetate by most definitions, especially as it is the most abundant species in the full community.

The lack of significant impacts of EKOs on other species was robust to using other methods of quantifying the impact of a species, such as using ANOVA (Supplementary Figure 1B). However, while we rarely observed multiple secondary impacts resulting from a single EKO, in some cases community biomass and structure did indeed change upon removing certain species (Supplementary Figure 2).

Overall, our ecological knockout experiments reveal a striking resilience of our communities to species removal, with a surprising lack of keystone species.

### Interaction matrix with a hierarchical structure decreases the likelihood of secondary impacts

In our synthetic marine communities, we observed a surprisingly low rate of secondary impacts over a wide range of environments. Is such a lack of keystone species what we should expect from a generic ecological community? To address this, we implemented a generalized Lotka-Volterra (gLV) model to simulate community assembly and EKOs of interacting species:

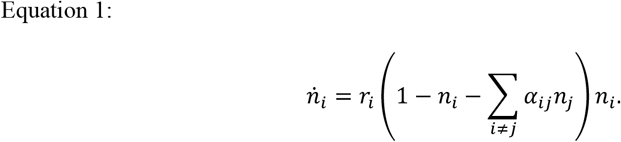

Here, *n*_*i*_ is population size that is normalized to each species’ carrying capacity. *r*_*i*_ is growth rate of species *i*, and *α*_*ij*_ is the interaction coefficient denoting impact from species *j* to species *i*. As a starting point, we simulated community assembly from 16-species pools with random interactions *α*_*ij*_ *∼ Uniform*(*μ* − *σ, μ* + *σ*), as we sweep over interaction strength *μ* and variation σ to represent a wide range of environment (Fig. 3A, Methods). This null model could capture variations in the number of co-existing species in the full communities in different resource environments, as the community diversity decreased with increasing mean and variance of the interaction strength (See discussion).

**Figure 3.**
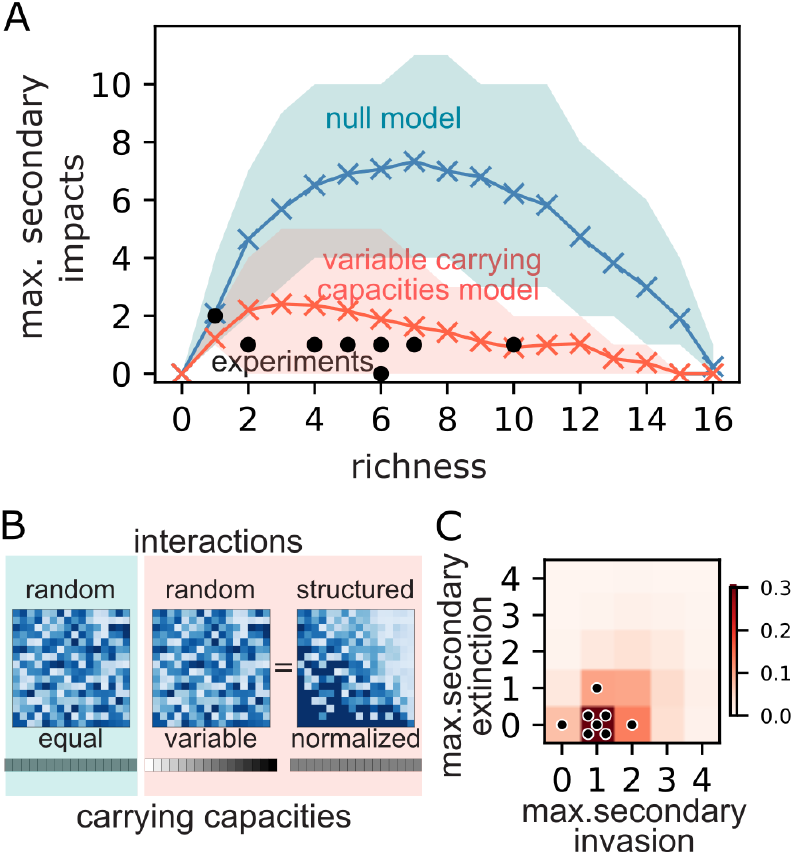
The generalized Lotka-Volterra model can recapitulate the low number of secondary impacts when structured interactions due to variation in species carrying capacities are included. **A**. Maximum secondary impacts for a given EKO in a simulation set / environment as a function of the richness of the full community. For the experimental data, each dot denotes the result of a certain environment (carbon source). For the simulations the mean over all simulations with a given full community richness appears in the plot, while the shaded regions denote the spread of the 10-90% of the data of each richness. The null model simulation is in blue; structured correlations model simulation is in red, and the experimental data is in black. **B**. Representative interaction matrices of the null model (left panel), in which the interactions are drawn independently and identically distributed (i.i.d) from a certain distribution (a uniform distribution is shown here), or a model incorporating species’ carrying capacities into the interaction matrix (middle and right - panels). The middle panel displays a model with interaction matrix drawn from uniform distribution and the introduction of variable carrying capacities, while the right panel displays an equal model in which the carrying capacities have been normalized and the interactions have been scaled accordingly (equation 3). Color denotes interaction strength with light blue equal to 0 and dark blue to 1.5. **C**. Density plot of the frequency of simulations/experiments resulting in the stated number of maximum secondary impacts, separately considering extinctions and invasions. In red is the color relevant for the frequency of this result in structured correlations simulation set. Each black dot represents an experimental environment.

To investigate the secondary impacts in our model, we calculated three metrics for each simulated species pool: (i) the number of co-existing species (richness) in the full community, (ii) the sum of all secondary impacts in each simulation species pool (total SI) and (iii) the maximum number of secondary impacts across all 16 EKOs (max SI). Our null model simulations showed a high number of max SI (4.4 ± 3.0), suggesting that we should naively expect most communities to have keystone species, as the removal of at least one species is expected to affect more than a quarter of the remaining species (Figure 2B). However, in contrast to the null model, our experimental observations revealed no keystone species. Specifically, among the eight environments, six had a max SI of one, one environment had a max SI of two, and another had none.

Moreover, most simulation sets of our null model resulted in a high total number of secondary impacts (18 ± 15), with 25% of individual EKOs leading to two or more secondary impacts. In contrast, we experimentally observed an average of 1.8 ± 1.0 secondary impacts per environment (total SIs), and only 1 out of 128 of EKOs resulting in two secondary impacts. Our gLV null model therefore massively over-predicts the number of secondary impacts relative to what we observe experimentally.

What could account for the discrepancy between our null model, which predicted numerous keystone species, and our experimental results, which demonstrated strong resilience to species removal? One possibility is that our null model neglected variation in carrying capacities between species as we considered interactions between populations normalized to their own carrying capacities. Indeed, our experimental species pool exhibited a wide range of carrying capacities in every environment (Fig. 1A), which was not explicitly considered in our null model. Therefore, as our revised model, we used a gLV model with variation in carrying capacities:

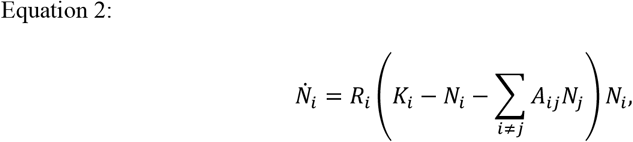

where *K*_*i*_ is the carrying capacity of species *i. N*_*i*_ is now the population size without normalization, and *R*_*i*_ and *A*_*ij*_ are converted accordingly (see SI Appendix). As before, we considered random interactions *A*_*ij*_ *∼ Uniform*(*μ* − σ, *μ* + σ), as we sweep over interaction strength *μ* and variation σ. Then we asked, unlike the null model case, this model with variation in *K*_*i*_ can let us to recapitulate the lack of keystone species.

We found that, as the variability of carrying capacities increased, secondary impacts became rarer in our simulations. In the case of a uniformly distributed carrying capacity between 0.1 and 1.9, we observed 1.9 ±1.3 max SI. Similarly, we observed a low total number of secondary impacts (3 ± 3) and 95% of individual EKOs in simulations led to 0 or 1 secondary impacts (Fig. 3A). Inclusion of variation in species carrying capacity in our model therefore recapitulated the small number of secondary impacts that we observed experimentally. Both with and without carrying capacity variation we observe a non-monotonic connection between secondary impacts and diversity, with intermediate richness leading to higher likelihood for secondary impacts (Fig. 3A).

How did the inclusion of carrying capacity variation reduce the secondary impacts? In the framework of gLV model, adding carrying capacities is equivalent to rescaling the interspecies interactions (Fig. 3B, Eq. 3, Appendix):

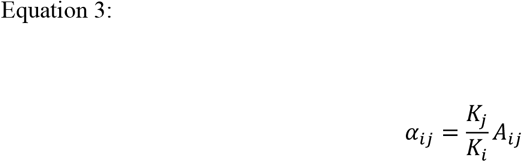

This indicates that, in terms of the normalized effective interactions *α*_*ij*_, species with a large carrying capacity therefore exert a stronger impact on species with a small carrying capacity. Relatedly, a species with a large carrying capacity is likely to be less impacted by all other species, which is reflected in the structure of the effective interactions (Fig. 3B right panel, Appendix). Therefore, our simulations suggest that structured interactions may prevent the emergence of keystone species.

It is natural to expect structured interactions in nature for many reasons including variation in carrying capacity and variation in growth rates coupled with mortality or dilution. However, it is counterintuitive that such structure would prevent, not promote, keystone species, given that a species with large carrying capacity might be expected to have a profound effect on a community, and removing the species might therefore lead to a cascade of secondary impacts as the community re-orders itself. To better understand how the emergence of keystone species depends on different types of structures, we took a closer look into the structure induced by carrying capacity. Carrying capacities introduce correlations to the pairwise interactions in a bi-directional manner: species with higher carrying capacities are likely to both strongly affect other species and to be weakly affected by others. We dissected the two directional effects by simulating EKOs with correlated interactions based on either affecter (⟨*α*_*ik*_*α*_*ik*_ *⟩* > 0) or affected (⟨*α*_*ik*_*α*_*ik*_*⟩*> 0) species:

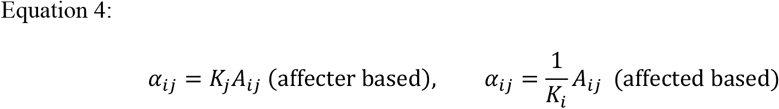

We found that only the affected-based correlation in interspecies interactions reduced secondary impacts, while the affecter-based correlation did not change the statistics of secondary impacts (Supplementary Figure 3). Perhaps in the presence of affected-based correlations, since a species is inhibited by a similar degree by all other species, removal of another species is unlikely to dramatically change the overall inhibition on it. This shows that structured interactions from carrying capacities or affected-based correlations can make communities resilient to removal of a species.

In addition to predicting that secondary impacts are rare, the variable carrying capacity gLV model also predicts that secondary invasions will be more common than secondary extinctions (with our base sampling parameters, simulations yielded 2.6±1.9 maximum secondary invasions across all 16 EKOs and 0.8 ±1.6 maximum secondary extinctions). Consistent with this prediction across the eight environments we observed 1.0 ±0.5 maximum secondary invasion across 16 EKOs and 0.1 ±0.3 maximum secondary extinctions (Fig. 3C). This contrasts with the null model in which the number of secondary invasions and extinctions are not only larger but also more balanced (supplementary figure 4).

### Inferred interspecies interactions display a hierarchical structure

We next sought to test whether the data from our experimental communities were consistent with structured interactions, predicted to reduce the prevalence of keystone species in gLV models. We did this in two complementary ways: (1) using statistical inference to fit the interaction strengths and infer any evidence of structure, and (2) using spent media experiments to analyze the impact of each species’ spent media on the growth of all other species.

First, we developed a statistical inference procedure to find the most likely matrix of interaction strengths that explained the observed species’ abundance data in different knockout experiments (Fig. 4A, see Methods). Briefly, our method used measurements of species’ individual carrying capacities (monoculture OD), relative abundances (from 16S amplicon-sequencing) as well as total community biomass (community OD), and assuming a Lotka-Volterra model, aimed to infer all interspecies interaction strengths that best explained the data. Since the number of fit parameters (interaction strengths) was larger than the number of data points (knocked out species), we used an appropriate penalty to avoid overfitting and used bootstrapping to infer a family of interaction matrices. For each matrix, we quantified the extent to which it had the structure expected from theory: namely the mean correlation between its columns (see Methods). We found that the distribution of inferred matrices had a much larger mean correlation than randomly shuffled versions of them (Fig. 4B). This was true across matrices inferred using all media tested (Supplementary Figure 5). Species abundances within communities were therefore consistent with the structured interactions as expected from theory.

**Figure 4.**
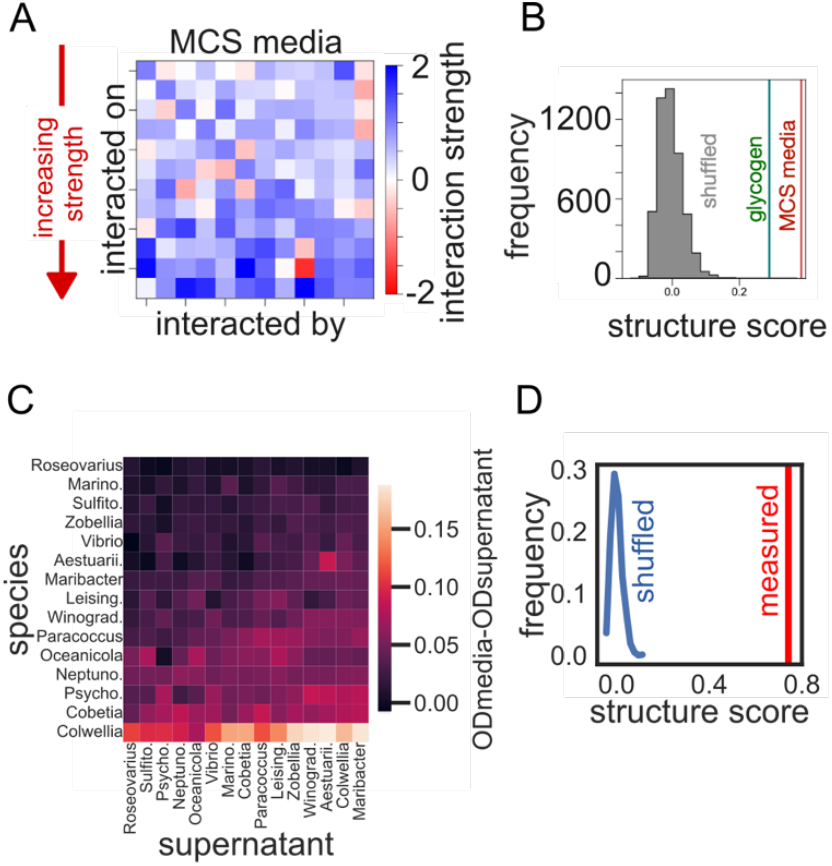
Inferred interaction matrix and spent media experiment suggest a hierarchical interaction structure between species. **A-B**. (left) Inferred interspecies interaction matrix from EKO community data in MCS media using our statistical approach. Species are ordered by increasing average interaction strength along rows (A). Structure score quantifying the mean correlation between columns of an interaction matrix (Methods) shown for matrices from MCS and glycogen media (B). Histogram shows a comparison with shuffled versions of these matrices. The matrices are much more structured than expected by chance, **C**. Measured supernatants impacts, calculated by the difference of OD at the end of 2 days of incubation on the supernatant and that of the original media. The rows are ordered by the least impacted species (mean impact from all species supernatants) to the most impacted. Similarly, columns are ordered by the least impacting supernatants to the most impacting. Reinekea sample was contaminated and removed from the analysis. **D**. Structure scores of the impacted species correlations. In red: the scores from the matrix appearing in C. In blue: a histogram of the scores of 1000 shuffles of the matrix entries.

As an independent way to infer interspecies interactions, we grew each species in the spent media of each other species. We quantified the relative growth of species *i* on the spent media of species *j*, compared with its growth in monoculture. This measure indicates the extent to which the environment created by species *j* promoted or hindered the growth of species *i*. We focused on the MCS media since all species were able to grow on it as monocultures (Fig. 1A), as well as it being the media with the highest richness of species (Fig. 1C). We found that the impacts a species experiences from other species supernatants were correlated and vice versa (Fig. 4C). This correlation was far higher than any correlation calculated on the shuffled matrix (Fig. 4D, Supplementary Figure 6 displays reproducibility), exemplifying the statistical significance of the correlation in the inferred interactions. We also observed similar correlations within a different species pool and environments by re-analyzing published data on pairwise interactions (Kehe et al^21^, Supplementary Figure 7.). Our spent media experiments therefore support the presence of structured interactions within the community, which provide a natural explanation for the absence of keystone species observed in our experiments.

## Discussion

In this work we used synthetic microcosms to systematically measure the frequency of secondary extinctions and invasions following the removal of each of the constituent species. To understand how secondary impacts depend on environmental resource complexity, we conducted our experiments using the same set of 16 species in eight distinct environments across a range of carbon sources with varying complexity. The assembled communities had varying levels of richness, ranging from one to twelve co-existing species. Strikingly, in none of the environments did we observe any species whose removal led to more than two secondary impacts, a finding made even more striking through comparison with a null model based on the gLV framework, which predicts that multiple secondary impacts should be common. Importantly, this lack of secondary impacts was observed even when removing species that are abundant within the original community, highlighting that our definition of keystone species is even less stringent than commonly used (where the impact of species removal is large compared with the abundance). Thus, we concluded that there were no keystone species in our microcosms in any environment.

We found that introducing correlations between the rows of the interaction matrix reduces the incidence of secondary impacts and thereby greatly diminishes the prevalence of keystone species in communities. The correlations introduced in the interaction matrix amount to creating a hierarchical structure of species interactions, rather than a random interaction network as is commonly assumed. Such correlations naturally arise when we consider the experimentally observed variation in carrying capacities or growth rates and mortality rates across different species in the gLV equations (Supplementary figure S8). Variation in any or all these quantities is expected, and results in an effective interspecies interaction matrix which is correlated or hierarchically structured (Fig. 3B), even though the underlying species-species interactions may be unstructured. The model also predicts that communities with intermediate richness tend to show the greatest number of secondary impacts (Fig. 3C, blue line), a pattern that can be understood via a simple theoretical argument based on random re-sampling of surviving species (Supplementary Discussion). However, secondary impacts in our experiment are too rare to confirm that such a pattern appears. By statistically inferring the effective interaction matrices using our community data, we indeed found that they possess such a hierarchical structure as expected from theory (Fig. 4). Additionally, when we measured pairwise interactions in spent media experiments, we observed a similar structure. An analysis of thousands of pairwise interactions measured in a droplet-based microfluidic chip^21^ show that these communities display similar structure as well (supplementary figure S7). Our results contribute to a recently growing understanding that soil and plant microbial communities might have a competitive hierarchy^22,23^. Overall, we believe that such structured interactions are abundant in microbial ecosystems and can provide resilience in the face of species extinctions and invasions.

Our results stand in contrast with several recent studies which performed similar EKO experiments to study microbiomes associated with specific environments such as plants^17–19^ and a “synthetic gut”^10,20^. These studies aimed to identify species with ‘keystone’ properties, using various metrics such as their influence on the host and changes in community structure. However, even when we relaxed our requirement for secondary invasions or extinctions and examined differences in species’ relative abundances between the full and EKO communities, our results generally support the lack of keystone species. Only one EKO showed significant changes in abundance in three species compared to the full community (Supplementary Figure 1B). When focusing on secondary impacts, two of these studies^10,20^ found a maximum of two secondary impacts, consistent with our findings. A third study^19^ identified an EKO with five secondary impacts, possibly due to their meticulous selection of the species pool. Future studies should explore how the degree and incidence of keystone species depend on the underlying species pool, the environment, and the degree of ecological and evolutionary adaptation of the species to the environment.

To summarize, our exploration of species removal across multiple environments shows that extinction/invasion cascades are rare compared to naïve theoretical expectations assuming uncorrelated interspecies interactions. Incorporating correlated interspecies interaction structure into account can help future theoretical studies reflect common ecological patterns, aid in the identification of keystone species for conservation efforts and enable the design of stable microbial communities in industry.

