## Supplemental Data for "Structured interactions explain the absence of keystone species in synthetic microcosms"

Cambridge 02139, USA.
^2^Institute of Biochemistry, Food Science and Nutrition. The Robert H. Smith Faculty of Agriculture, Food and Environment. The Hebrew University of Jerusalem. 229 Herzl Street, Rehovot 7610001, Israel.

^3^International Centre for Theoretical Sciences, Tata Institute of Fundamental Research, Bengaluru 560089, India

*Corresponding author

Conversion between gLV equations with and without carrying capacity variations

In the main text, we discuss two ways of writing generalized Lokta-Volterra models (eq. 1, eq. 2). Here, we will explicitly calculate the conversion from Equation 2, which models gLV dynamics of populations with varying carrying capacities, to Equation 1, which models gLV dynamics of normalized populations with respect to each species’ carrying capacities.

Let us start from Equation 2:

$$\dot{N}_{i}=R_{i}\left( K_{i}-N_{i}-\sum_{i\neq j} A_{ij}N_{j} \right)N_{i}.$$

We now replace the population sizes $N_{i}$ with normalized population sizes $n_{i}=N_{i}/K_{i}$:

$$K_{i}\dot{n}_{i}=R_{i}\left( K_{i}-K_{i}n_{i}-\sum_{i\neq j} A_{ij}{K_{j}n}_{j} \right){K_{i}n}_{i},$$

which can be simplified to

$$\dot{n}_{i}=R_{i}K_{i}\left( 1-n_{i}-\sum_{i\neq j} \frac{K_{j}}{K_{i}} A_{ij} n_{j} \right)n_{i}.$$

This is equivalent to Equation 1 with $r_{i}=R_{i}K_{i}$ and $\alpha_{ij}=\frac{K_{j}}{K_{i}} A_{ij}$.

Calculation of secondary impacts in the case of random assembly

In the main text, we discuss how the number of secondary impacts in simple Lotka-Volterra model reaches a null model prediction, i.e., the case of random assembly. Here we discuss how the number of secondary impacts can be calculated for the case of random assembly.

For the case of random assembly, we assume that the community assembly after an ecological knockout becomes completely independent of the community before the knockout, except that the average probability for survival over all species remains the same. In other words, this is a situation in which the survival of each species before the knockout does not provide any information on whether it can survive after the knockout. Under such assumption, we can calculate the expected number of secondary impacts in the following way.

Let us consider a situation in which the species pool size is $S_{0}$ and the chance of survival of a species in the given environment is *S*/$S_{0}$. In a specific scenario we expect to have *S* species survive in the full community. Now let us calculate the expected number of secondary impacts by considering two different situations.

First, let us consider the case in which a EKO was originally surviving, which has a chance of $S/S_{0}$. Then there are two ways that secondary impacts can occur: an originally survived species going extinct, and an originally extinct species surviving. In the former case, $S-1$ other species which originally survived, there is $1-S/S_{0}$ chance that they go extinct. Similarly in the latter case, for $S_{0}-S$ species which originally went extinct, there is $S/S_{0}$ chance that they survive. In the end, in the case in which an EKO species was originally surviving, the expected number of secondary impacts is $\frac{S}{S_{0}}\left( 1-\frac{S}{S_{0}} \right)\left( \frac{2S-1}{S_{0}} \right)$ .

Next, let us consider the case in which a KO’ed species was originally going extinct, which has a chance of $(1-\frac{S}{S_{0}})$. Then for $S$ species which originally survived, there is $1-S/S_{0}$ chance that they go extinct. Similarly, for $S_{0}-S-1$ species which originally went extinct, there is $S/S_{0}$ chance that they survive. In the end, in the case in which a KO’ed species was originally extinct, the expected number of secondary impacts is $\frac{S}{S_{0}}\left( 1-\frac{S}{S_{0}} \right)\left( \frac{2S_{0}-2S-1}{s_{0}} \right)$.

Overall, the expected number of secondary impacts in the null model of random assembly is given by

$$2\frac{S}{S_{0}}\left( 1-\frac{S}{S_{0}} \right)(1-\frac{1}{S_{0}})$$

### Supplementary methods

**Minimal media**

Seawater (342.25mM NaCl, 14.75mM MgCl_2_-6H_2_O,1mM CaCl_2_-2H_2_O, 6.75 KCl) mixed with trace minerals and vitamins (2.1mg/L FeSO_4_ * 7H_2_O, 0.03 mg/L H_3_BO_3,_ 0.1 mg/LMnCl_2_ * 4H_2_O, 0.19 mg/L CoCl_2_ * 6H_2_O, 0.24 mg/L NiCl_2_ * 6H_2_O, 0.2 mg/L CuCl_2_* 2H_2_O, 0.144 mg/L ZnSO_4_ * 7H_2_O,0.036 mg/L Na_2_MoO_4_ * 2H_2_O, 0.025 mg/L NaVO_3_, 0.025 mg/L NaWO_4_ 2H_2_O, 0.006 mg/L Na_2_SeO_3_ 5H_2_O, 0.1 mg/L Riboflavin, 0.03mg/L D-Biotin, 0.1 mg/L Thiamine hydrochloride, 0.1 mg/L L-ascorbic acid, 0.1 mg/L Ca-d- pantothenate, 0.1 mg/L Folate, 0.1 mg/L Nicotinate, 0.1 mg/L 4-aminobenzoic acid, 0.1 mg/L Pyridoxine HCl, 0.1 mg/L Lipoic acid, 0.1 mg/L NAD, 0.1 mg/L Thiamin pyrophosphate, 0.01 mg/L) Cyanocobalamin that were donated by the Cordero lab. Nitrogen source: 0.01 M ammonium chloride, phosphorus source: 1 mM phosphate dibasic, sulfur source: 1 mM sodium sulfate. Different carbon sources were added as a function of the experiment. The carbon sources used are listed in Table S2 below, as well as their MW if applicable.

Table S1 - Strains

| Isolate | Genus | Family | order | class |
| --- | --- | --- | --- | --- |
| I2R16 | *Psychosphaera* | *Pseudoalteromonadaceae* | *Alteromonadales* | *Gammaproteobacteria* |
| 3B05 | *Neptunomonas* | *Oceanospirillaceae* | *Oceanospirillales* | *Gammaproteobacteria* |
| G2M07 | *Reinekea* | *Oceanospirillaceae* | *Oceanospirillales* | *Gammaproteobacteria* |
| C2R09 | *Paracoccus* | *Rhodobacteraceae* | *Rhodobacterales* | *Alphaproteobacteria* |
| C3M06 | *Oceanicola* | *Rhodobacteraceae* | *Rhodobacterales* | *Alphaproteobacteria* |
| E3R09 | *Winogradskyella* | *Flavobacteriaceae* | *Flavobacteriales* | *Flavobacteriia* |
| E3R11 | *Vibrio* | *Vibrionaceae* | *Vibrionales* | *Gammaproteobacteria* |
| A2M03 | *Flavobacteriaceae* | *Flavobacteriaceae* | *Flavobacteriales* | *Flavobacteriia* |
| C3M10 | *Leisingera* | *Rhodobacteraceae* | *Rhodobacterales* | *Alphaproteobacteria* |
| B3M08 | *Sulfitobacter* | *Rhodobacteraceae* | *Rhodobacterales* | *Alphaproteobacteria* |
| C2M11 | *Colwellia* | *Colwelliaceae* | *Alteromonadales* | *Gammaproteobacteria* |
| B2M13 | *Cobetia* | *Halomonadaceae* | *Oceanospirillales* | *Gammaproteobacteria* |
| A3R04 | *Aestuariibacter* | *Alteromonadaceae* | *Alteromonadales* | *Gammaproteobacteria* |
| C3R19 | *Maribacter* | *Flavobacteriaceae* | *Flavobacteriales* | *Flavobacteriia* |
| D2R04 | *Roseovarius* | *Rhodobacteraceae* | *Rhodobacterales* | *Alphaproteobacteria* |
| F3R08 | *Marinobacter* | *Alteromonadaceae* | *Alteromonadales* | *Gammaproteobacteria* |

Table S2 Carbon sources

| Carbon Source | Final M | Final carbon M | Final % | MW (g/mole) |
| --- | --- | --- | --- | --- |
| sodium acetate  (Sigma 791741) | 0.06 | 0.12 | 0.49 | 82.03 |
| Glucose  (Sigma G8270) | 0.02 | 0.12 | 0.36 | 180.16 |
| GlcNaC  (Sigma A4106) | 0.015 | 0.12 | 0.33 | 221.21 |
| Sucrose  (Macron 279110) | 0.01 | 0.12 | 0.34 | 342.3 |
| D-Raffinose pentahydrate (Amresco J392) | 0.0067 | 0.12 | 0.4 | 594.51 |
| sodium alginate  (Sigma W201502) | 0.0058 | 0.035 | 0.125 | 12000-14000 |
| glycogen  (Sigma [G8751](https://www.sigmaaldrich.com/IL/en/product/sigma/g8751)) | 0.0019 | 0.045 | 0.125 | 270-3.5x10^6^ (Personal correspondence with Sigma representative.) |
| MB diluted 1:5  (BD difco 279110) | NA | NA | 0.0748 | NA |

All components mentioned here were weighted, mixed (titrated and stir-heated if needed) and filtered through a 0.2uM PES filter.

**DNA extraction and 16S-amplicon sequencing**

Samples were defrosted, 200ul were moved to 96 U-bottom well plates (Corning #353077), and centrifuged at 3,220rcf for 3 minutes to pellet cells. Cells were washed with sterile water, pelleted again, and then supernatants were aspirated. Cells were resuspended in 25ul of TES buffer (10 mM Tris-HCl, 1 mM EDTA, 100 mM NaCl). To lyse the cells, 250U/ul of ReadyLyse (Lucigen R1804M) were added to the samples, followed by overnight shaking at room temperature. Samples were then centrifuged for 5 minutes at 3,220rcf and supernatant was stored at -20C till shipment to sequencing. Prior to freezing, 1ul of each sample were taken to measure DNA concentration using Quant-it PicoGreen (ThermoFisher, P7589). Samples were sent to Aragonne National Laboratory where 16S-amplicon libraries were prepared with the 515F-860R primer set and sequenced on an Illumina MiSeq machine with a 2x151bp run.

Amplicon sequence variants (ASVs) were obtained using the R package DADA2. Filtering and trimming carried with the parameters: truncLen=c(150,150), trimLeft = 10, maxN=0, maxEE=c(2,2) and truncQ=2.

Taxonomic identities were assigned to ASVs using the SILVA version 138 database train set. R4.1.3 was used.

To match the assigned ASVs to any of the 16 species used in the experiment, BLAST (blastn version 2.13.0+) was run locally against a database containing the 16S sequences obtained from the 16 species. An ASV was assigned if there were no more than two mismatches between the query and the hit from the database. ASVs without a match were grouped as “other”. 15 samples with ‘other’ assigned to more than 10% of the reads were removed from the analysis. Also 1 sample with less than 1000 reads was also removed from the analysis.

This resulted in a dataset of 312 samples, with a mean of 23,361±10,147 reads per sample with an average frequency of less than 0.77%± 1.62% of reads assigned as ‘other’.

**Model and simulation**

For carrying capacity, we use unit $K_{i}=(1, \ldots., 1)$ for the null model and an arithmetic sequence $K_{i}=(0.1, 0.22, \ldots, 1.78, 1.9)$ for the carrying capacity model.

For interspecies interaction, we sweep over two parameters $\left( A_{min},A_{max} \right)$ that determine the strength and variation of interaction matrix $A_{ij}$ with steps size of 0.075. In other words, the distributions we use for sampling $A_{ij}$ are $Uniform\left( 0,0 \right),$ $Uniform\left( 0, 0.075 \right)$, $Uniform\left( 0.075, 0.075 \right)$, …, $Uniform\left( 1.5,1.5 \right)$.For each parameter set $\left( A_{min},A_{max} \right)$, we draw 100 communities $A_{ij}$ with 16 species. In this way, we sample total of 19000 communities for each model.

For each community (i.e. for each choice of $A_{ij}$), we numerically solve dynamics with initial conditions $N^{full}\left( t=0 \right)=(\frac{1}{16},\ldots,\frac{1}{16})$ for the full community simulation and $N^{EKO_{i}}\left( t=0 \right)=0 for species i, \frac{1}{16} for all others$ for 16 EKO simulations. We use numerical solver Tsit5 with automatic switching based on Rosenbrock method. This is done with package Differential Equations in Julia (v 1.9). Any species with population fraction < 0.01% at time $t=1000$ after the onset of simulation is considered extinct.

**Statistically inferring interactions from data**

To infer the interaction matrices from the community abundance data, we took a maximum likelihood approach. Our method is inspired by recent approaches to infer interactions but operates in a different regime, one where coexistence is not common and ecological knock-out experiments do not provide enough data for unambiguous inference^27,28^. For each species, we had measurements of their carrying capacities *K* from monoculture experiments, as well as relative abundances in knockout communities. For each knockout community, we also had measurements of the community biomass in the form of its total optical density (OD), which we used to convert each species’ relative abundance to absolute abundance in units of OD. For each species gamma, we thus had a measured abundance N_obs, as well as a predicted abundance N_pred under a generalized Lotka-Volterra model (gLV) with an interaction matrix A to be determined. Generally, for any community with a set of surviving species (indicated by an asterisk *), the gLV prediction for the abundance of species gamma was given by:

$$N_{obs,\gamma}= \sum_{\delta=1}^{S} A_{\gamma\delta}^{-1}K_{\delta}$$

Assuming log-normally distributed errors (a reasonable assumption for microbiome abundance data^29^), we thus minimized the following cost function, which included the following two elements: (1) the total likelihood of observing a given set of species abundances in communities, as well as (2) a Ridge regularization penalty to avoid overfitting and ensure that the inferred matrices did not have arbitrarily large interaction strengths:

$$A_{opt}=\arg\min_{A} [\sum_{i=1}^{N} \sum_{\gamma=1}^{S} \left( \log N_{obs,\gamma}-\log N_{pred,\gamma} \right)^{2}-\lambda\left| \left| A \right| \right|^{2}]$$

We used the matrix *A_opt_* that minimized the above cost function as the inferred matrix for the given community data. Note that here, we used data from all knockout communities (represented by the letter *i*) for a given growth medium and over all species present in that community (represented by the symbol γ), since we would generally expect different media to yield different interaction matrices for different communities.

**Growth abilities on different species supernatants**

Starter cultures were prepared twice, once for the supernatants and once for the species, in the following way: colonies from all 16 isolates streaked from frozen stock on MB-agar plates were picked, placed into culture tubes with 3ml MB, and grown at room temperature with shaking at 300rpm for two days.

For the supernatant cultures: each culture was diluted 1:40 in fresh MCS media for a total of 22.5ml in 50ml Falcon tubes. These cultures were grown at room temperature for 2 days, shaking. All cultures but that of *Reinekea* were filtered through 0.2um filter (ThermoFisher VWR#82030-938). *Reinekea* was filtered through 0.1um filter (ThermoFisher #565-0010), since this species can pass the 0.2um filter. These supernatants were diluted 1:1 with fresh MCS media.

For the growth experiment starter cultures from each species were diluted 1:40 in 400ul of either supernatant or MCS media in triplicates. This was done in 1ml deep 96-well plates (Eppendorf # 951033006). These plates were shaken for 2 days at RT. Then 100ul of each sample, as well as background media were taken to have OD_600_ measured in a multiplate reader.

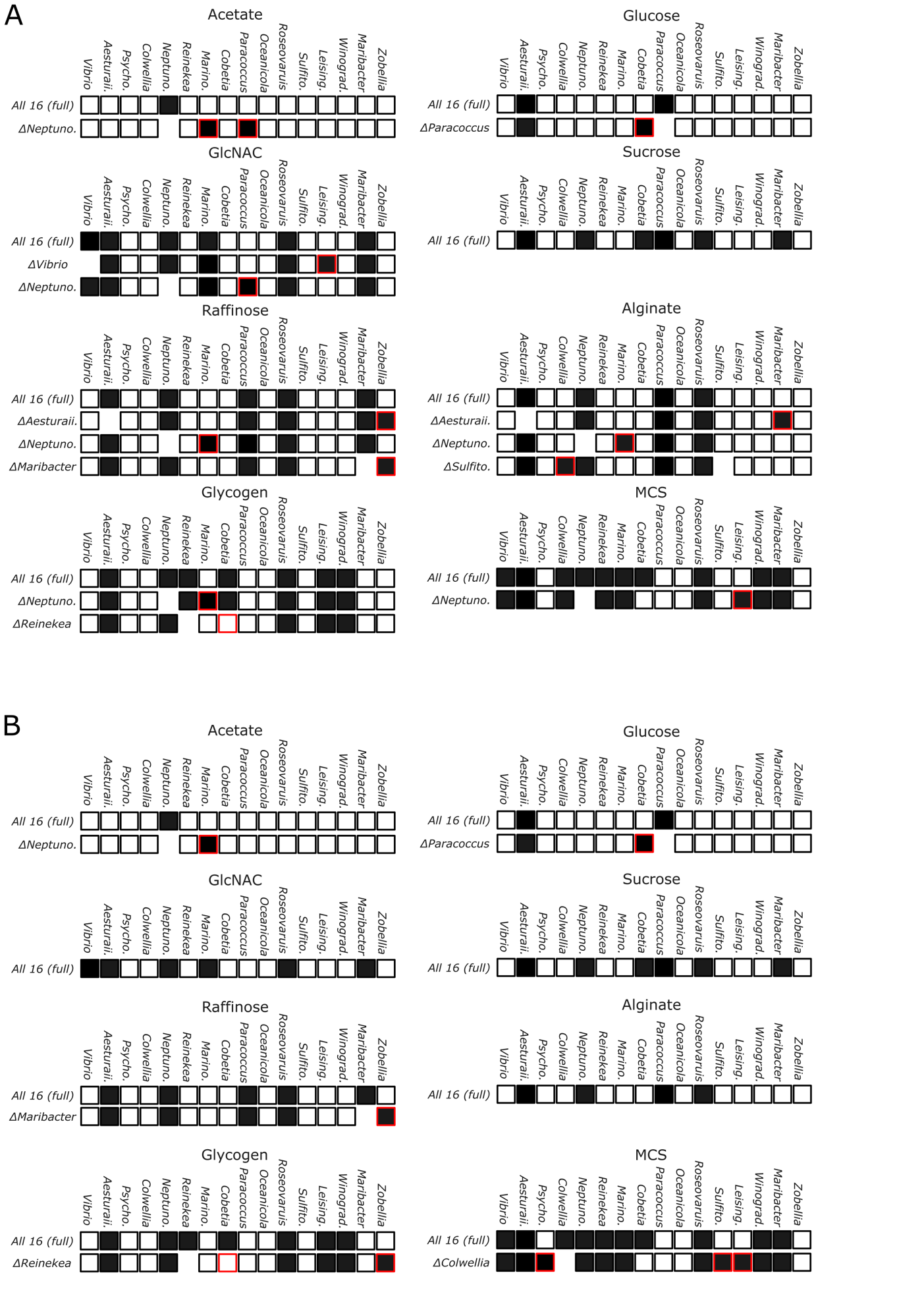

Supplementary Figure 1. Lack of keystone species is robust to the method used. **A.** Secondary impact analysis of all media, as explained in Figure 2B. Here only EKO communities that lead to secondary impacts are displayed, for all media. **B**. Secondary impacts analysis similar to the one presented on Figure 2. Here secondary impacts were inferred using ANOVA analysis following CLR conversion of the read counts. In this analysis we have first identified for each media and species in which EKOs this species is significantly different. Then for the significantly impacted species we have searched for a significant EKOs that is different from the full community. This was done using the statsmodels module in python.

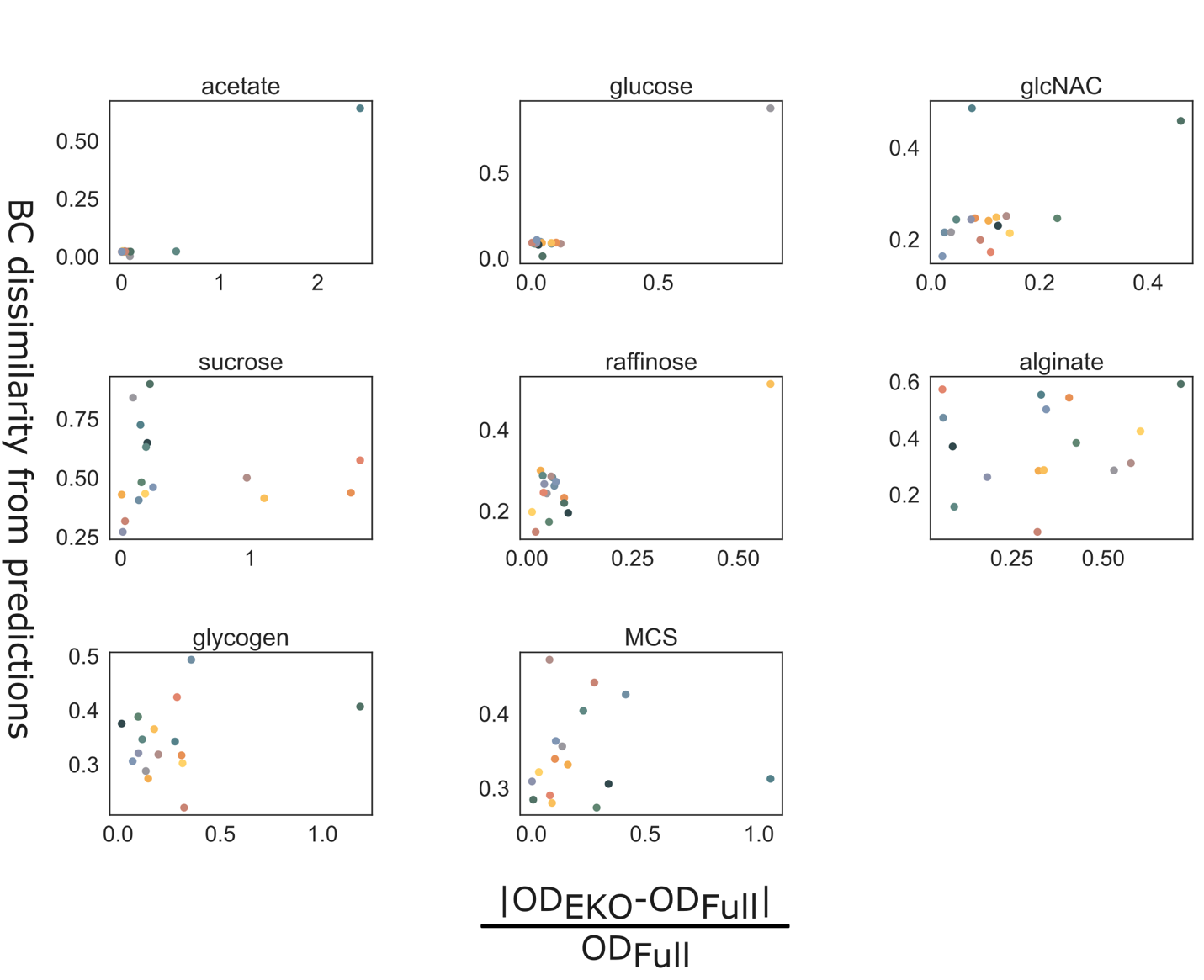

Supplementary Figure 2 Community shift vs. OD changes. For each EKO, its Bray-Curtis dissimilarity was calculated based on its prediction (normalized ASV frequencies after removing the EKO species reads from the full community). This was plotted against the normalized difference in OD of the EKO community compared to the full community. The two distance measures are correlated, with Spearman's rho = 0.6 and a p-value of 4.3 x 10^-14. Each dot represents the mean distances of all EKO replicates from the full community triplicates.

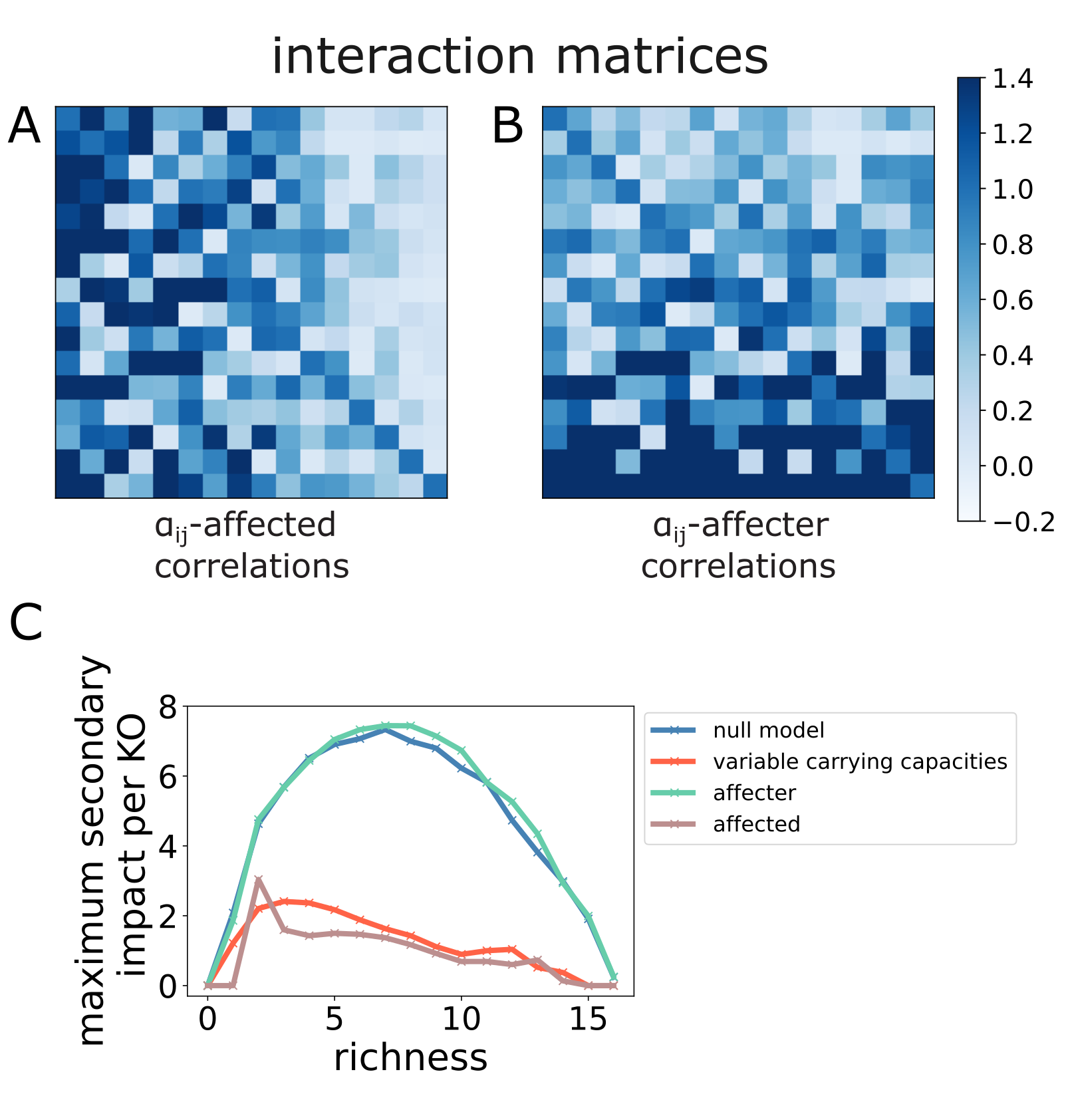

Supplementary Figure 3. Correlation of the affected interactions are the main contribution to the decrease in secondary impacts. **A-B.** Representative interaction matrices with correlations based on the affected species (A) and on the affecter species (B). The base interactions $A_{ij}$ are drawn independently and identically distributed (i.i.d) from a certain distribution (a uniform distribution is shown here), and then the effects from carrying capacity $a_{ij}=\frac{K_{j}}{K_{i}}A_{ij}$ is applied only across different rows (A: $a_{ij}=\frac{1}{K_{i}}A_{ij}$) or across different columns (B: $a_{ij}={K_{j}A}_{ij}$) of interaction matrices. **C**. Maximum secondary impacts per EKO as a function of the richness of the full community. 4 curves are shown: affected-based correlations, affecter-based correlations, carrying capacity-induced correlations, and no correlations (the null model). We find that both affected-based correlations and carrying capacity-induced correlations lead to decrease in secondary impacts, while affecter-based correlation does not change secondary impacts compared to the null model.

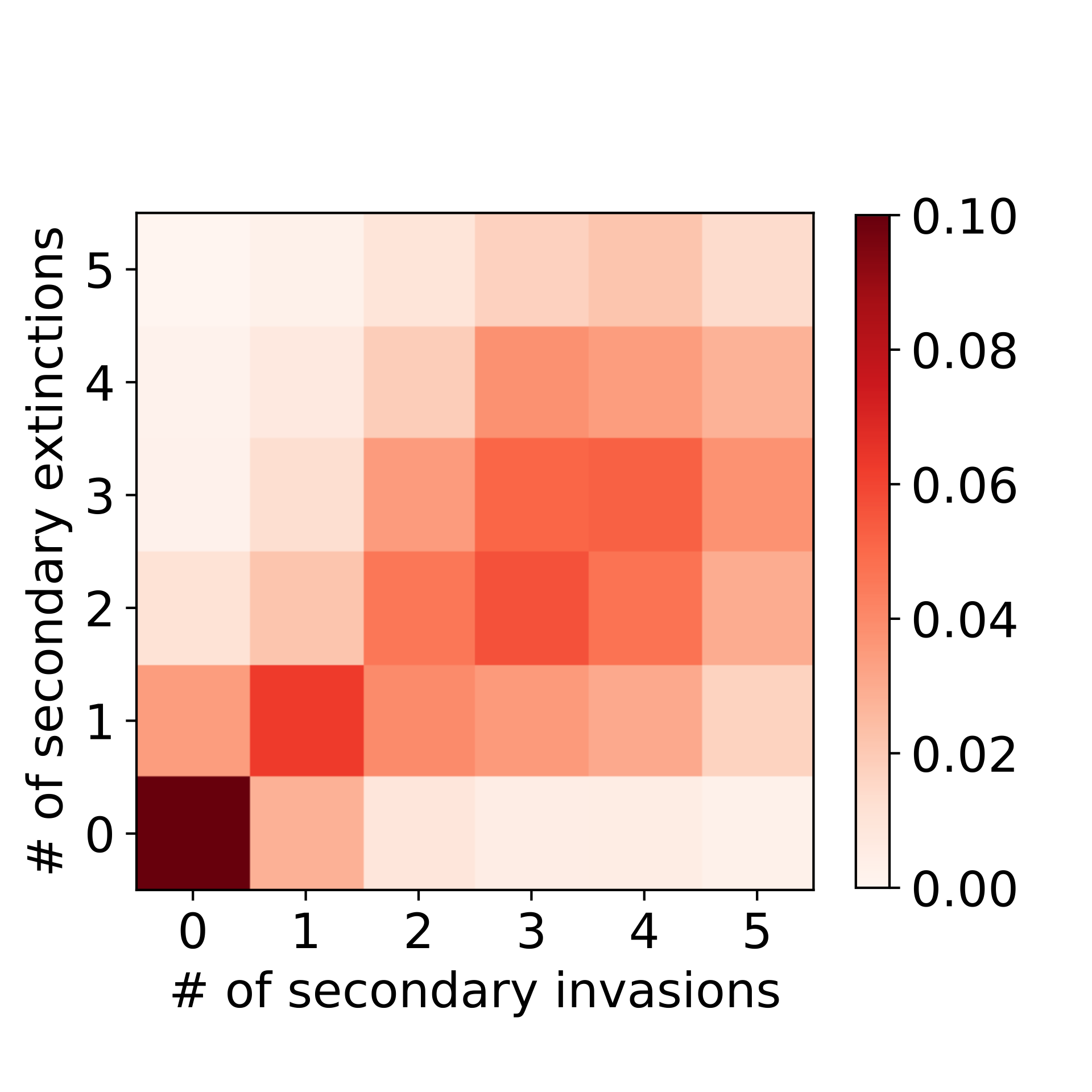

Supplementary Figure 4. Null model has a similar likelihood of secondary invasions and extinctions. Density plot of the frequency of null model simulations resulting in the stated number of secondary extinctions and invasions. The data is distributed along the line $x=y$, unlike in the case of carrying capacity simulations (Fig. 3C).

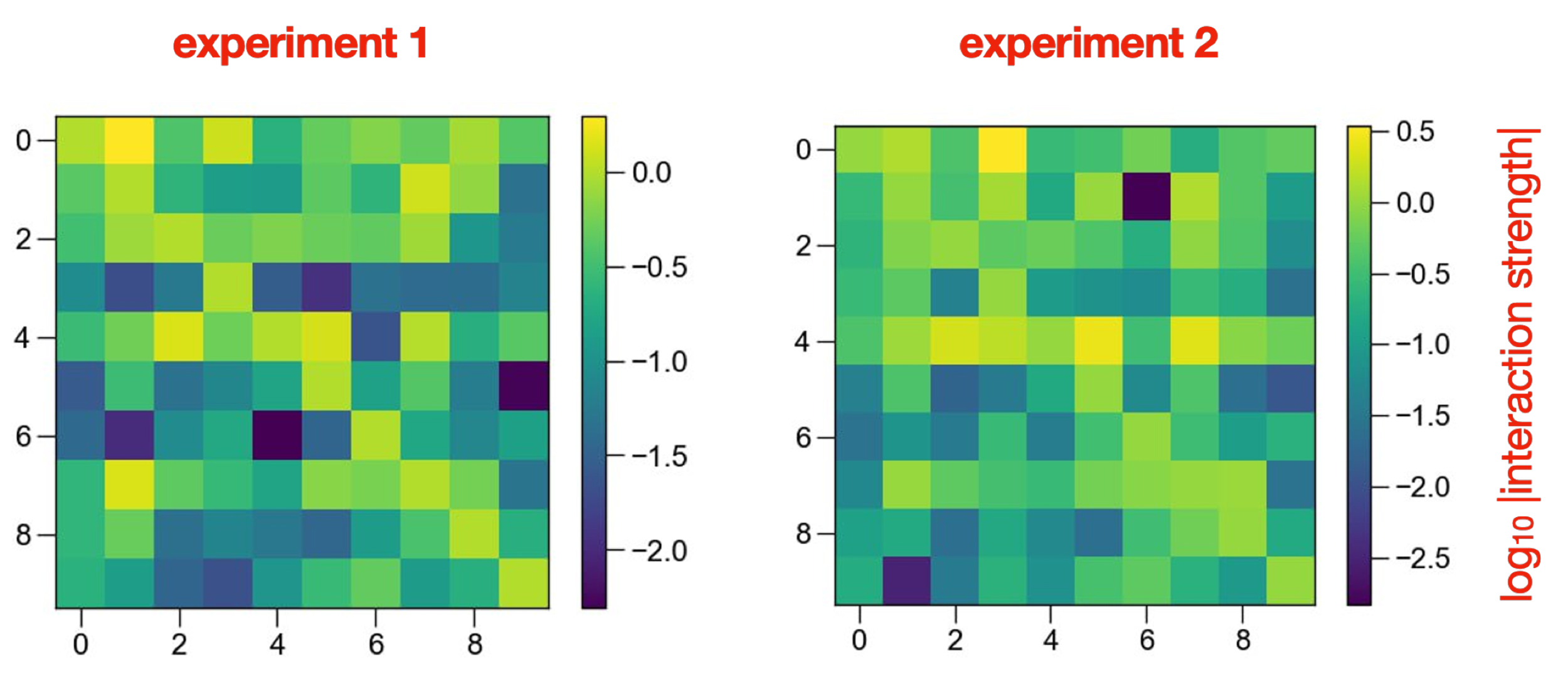

Supplementary Figure 5. Inferred interaction matrices for multiple environments. Interaction strength matrices for the 9 species that coexist in the glycogen communities inferred using data from two independent experimental replicates (left and right). The color bars represent the log of the magnitude of the interaction strength between pairs of species. The two matrices are strongly correlated (R = 0.88; P < 10^-3^).

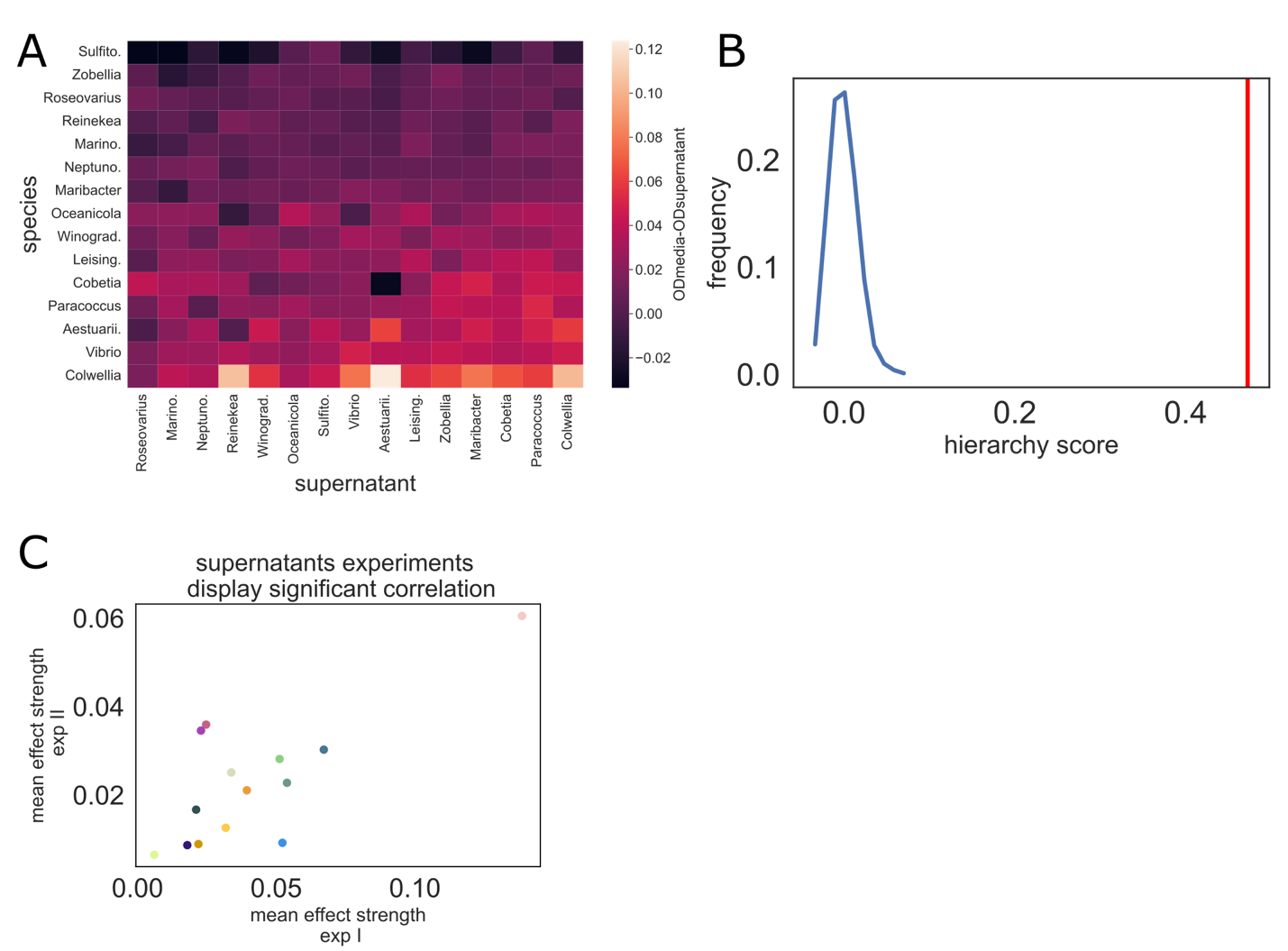

Supplementary Figure 6 Repetition of the spent media experiment show similar hierarchical structure. A. measured supernatants impacts. The rows are ordered by the least impacted species (mean impacts from all supernatants) to the most impacted. Similarly, columns are ordered by the least impacting supernatant to the most impacting. B. Hierarchical scores of the species correlations. In red the score of the measured matrix in A. In blue a histogram of the score of 1000 shuffles of the matrix entries. C. a scatter plot of the mean of each species impacts from the 2 experiments (the one from Figure 4 and this one). The Pearson correlation between them is significant ( ρ= 0.755, p-value=0.0018).

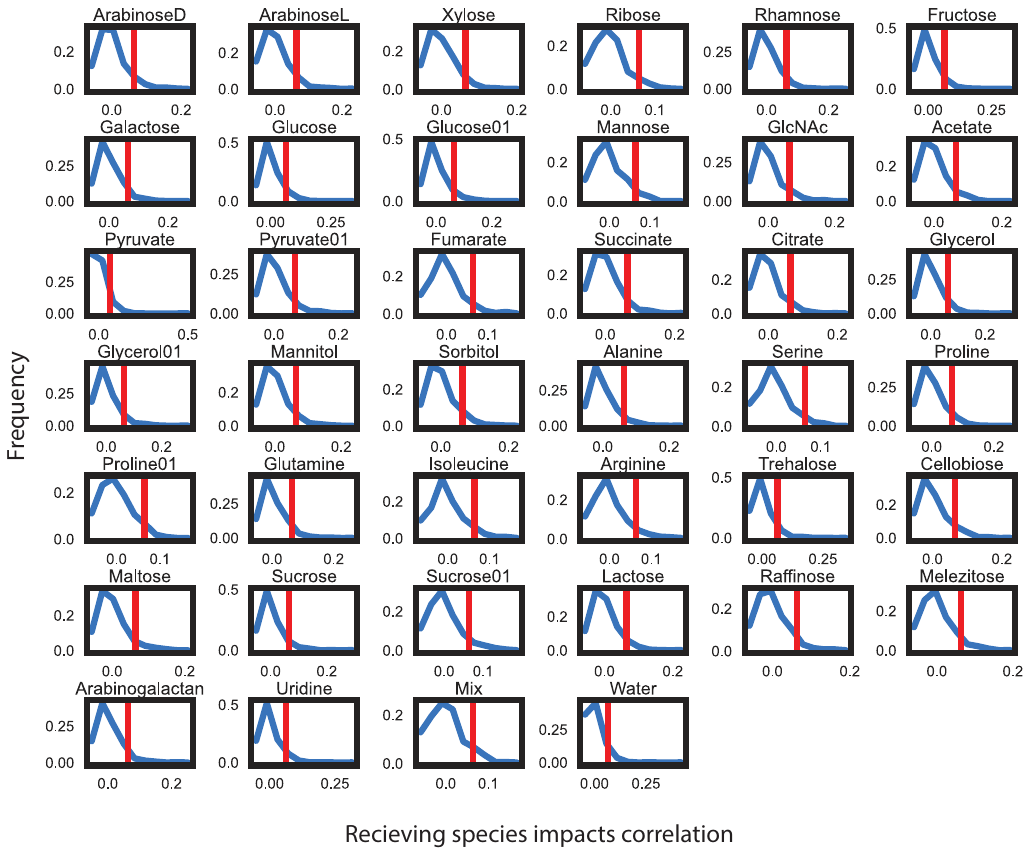

Supplementary Figure 7. Pairwise interactions measured using the K-chip display columns correlations. Scores of the impacted species correlations over 40 different carbon sources tested in the K-Chip^21^. In red- the score of the measured interaction matrix. In blue a histogram of the score of 1000 shuffles of the matrix entries.

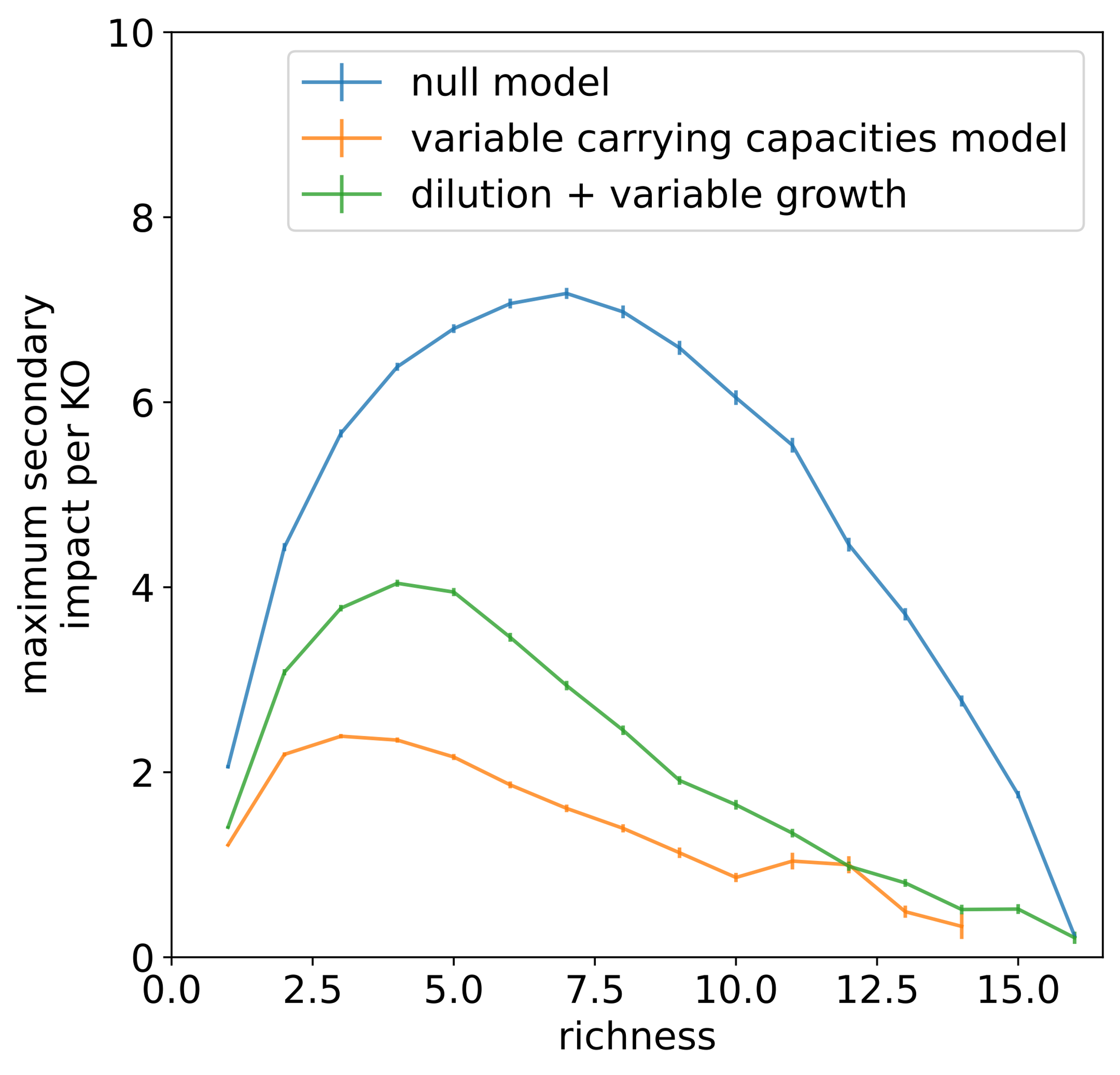

Supplementary Figure 8. Variable growth rate and dilution can reduce secondary impacts. Under dilution+variable growth condition, growth rates are evenly distributed across 16 species from 0.5 to 1.5 and a universal mortality of 0.4 is applied for all species. The plot shows results from 19000 communities as in the main text. We find that combining variation in growth rates and dilution can reduce the secondary impacts. This is because dilution rate structures the effective interaction matrix according to growth rate hierarchy^30^.
